# The phosphoinositide PI5P impairs mitochondrial function through endosome-mitochondria proximity

**DOI:** 10.1101/2025.09.22.677701

**Authors:** Helene Tronchere, Sourour Ben Harzallah, Mathieu Cinato, Valentin Antoszewski, Vanessa Soldan, Stéphanie Balor, Oksana Kunduzova, Frank Lezoualc’h, Frederic Boal

## Abstract

Alterations of mitochondrial structure and bioenergetic dysfunction are hallmarks of metabolic stress and in the progression of heart failure (HF). However, the impact of lipid signalling, particularly the dynamics of phosphoinositides is not fully characterized. Here, we investigate the effect of phosphatidylinositol 5-phosphate (PI5P) on mitochondrial structure and function in cardiomyocytes. We show that exogenous PI5P induces mitochondrial fragmentation, loss of mitochondrial membrane potential, increased ROS production, and impaired mitochondrial respiration. Using a PI5P biosensor, we identify a significant mitochondrial PI5P pool, whose abundance increases upon metabolic stress induced by 2-deoxy-D-glucose exposure, most likely in a PIKfyve-dependent manner. Furthermore, we demonstrate that PIKfyve-positive endolysosomes localize in close proximity to mitochondria in an actin-dependent manner, suggesting a spatial and functional coupling between these two organelles. Overall, our findings reveal a novel mechanism by which PI5P contributes to mitochondrial dysfunction during metabolic stress, potentially linking endolysosomal signalling to mitochondrial metabolism and cardiomyocyte bioenergetic status.

## Introduction

Phosphoinositides (PIs) are a family of phospholipids that play key roles in intracellular signalling, membrane trafficking, and organelle identity ^1^. Despite its low abundance, phosphatidylinositol 5-phosphate (PI5P) has emerged as a signalling lipid involved in various cellular processes, including nuclear gene expression ^2^, Akt signalling pathway ^3^ and endosomal maturation ^4^. PI5P is recognized as a stress lipid, as its synthesis is triggered by various cellular stresses, including thrombin stimulation in human platelets ^5^, T-cell activation ^6^, oncogenic malignancy ^7^, osmotic shock ^8^ and UV irradiation ^2^. PI5P is produced by the ubiquitous lipid 5-kinase PIKfyve ^9,10^, either directly from phosphatidylinositol or indirectly via the subsequent action of the 3-phosphatases of the myotubularin family on PI(3,5)P_2_ ^11,12^. PIKfyve is a large multi-domain ubiquitous protein. It contains a conserved Fab1p/YOTB/Vac1p/EEA1 (FYVE) domain that binds PI3P and targets the protein to endolysosomal membranes ^13,14^, a C-terminal phosphoinositide kinase domain, and several coiled-coil and cysteine-rich regions likely involved in protein-protein interactions ^15^. PIKfyve forms a ternary complex with the scaffolding protein Vac14 and Fig4, a Sac3-domain-containing phosphoinositide phosphatase and this complex is required for PIKfyve enzymatic activity ^16–18^. Functionally, PIKfyve is essential for endosomal membrane dynamics, lysosome homeostasis, and autophagic flux ^19–23^, as well as retrograde trafficking ^24,25^. Pharmacological inhibition of PIKfyve has emerged as a valuable tool to investigate its cellular functions. Among several described inhibitors, Apilimod, also known as STA-5326 (referred as STA in this study) has since been identified as a potent and selective PIKfyve inhibitor, efficiently blocking the production of PI(3,5)P₂ and PI5P.

The heart is a high-energy demanding organ, consuming an average of ∼6 kg of ATP per day ^26^. Therefore, cardiac performance strongly depends on continuous mitochondrial ATP production, and even transient disturbances in energy supply can rapidly impair contractile function, leading to heart failure (HF), a major cause of mortality worldwide. Under conditions of chronic pressure overload, increased workload, or nutrient limitation, cardiomyocytes experience metabolic stress, disrupting mitochondrial bioenergetics and contributing to the progression of HF.

In this context, our previous work has demonstrated that PIKfyve plays an essential role in maintaining mitochondrial structural integrity under hypoxic or metabolic stresses in cardiomyocytes ^27^. Specifically, we found that pharmacological inhibition of PIKfyve reduced stress-induced ROS production and prevented apoptotic cell death— two key determinants of HF progression. These findings highlighted a potential connection between PIKfyve-dependent lipid signalling and mitochondrial adaptation to metabolic stress.

In the present study, we show that an increase in PI5P levels leads to elevated ROS production, reduced mitochondrial respiration and affects mitochondrial structure in cardiomyocytes. In addition to a vesicular PIKfyve-PI5P pool, we demonstrate the existence of a significant PIKfyve-dependent PI5P pool localized at the mitochondria, that contributes to mitochondrial dysfunction. Under metabolic stress, mitochondrial PI5P level and the proximity of PIKfyve-positive endolysosomes to mitochondria increase. These findings uncover an unrecognized role of PIKfyve in regulating mitochondrial homeostasis in energy-demanding cardiomyocytes via spatially defined pools of PI5P.

## Results

### Increased levels of PI5P triggers mitochondrial dysfunction

We previously demonstrated that pharmacological inhibition of PIKfyve prevented mitochondrial ROS production and mitochondrial structure alterations induced by hypoxic or metabolic stress in cardiomyocytes, suggesting that PI5P production could alter mitochondrial structure and function. To investigate the effect of an artificial increase in PI5P levels on mitochondria, we first resorted to the expression of the bacterial phosphatase IpgD which hydrolyses PI(4,5)P_2_ to generate high amount of intracellular PI5P ^28^. We found that expression of IpgD induced a significant fragmentation of the mitochondrial network, while the catalytically inactive mutant IpgD^C438S^ did not induce such fragmentation (Figure 1A). Conversely, expression of the yeast phosphatase Inp54p, which catalyzes the conversion of PI(4,5)P_2_ to PI4P, had no effect on mitochondrial structure, indicating that neither the drop in PI(4,5)P_2_ levels not the rise in PI4P were responsible for the observed mitochondrial fragmentation (Figure 1A). To further confirm these results, we next used short-chain cell-permeant PI5P, which has been shown to recapitulate some effects of IpgD expression ^3,4,29–31^. As shown in Figure 1B, treatment with exogenous PI5P induced a mitochondrial fragmentation, similarly to IpgD expression, while treatment with PI4P had no effect. This was accompanied by an increase in mitochondrial ROS production (mtROS, Figure 1C) and a reduction in mitochondrial membrane potential (mtΔΨ), as highlighted by the TMRE staining (Figure 1C). We then sought to investigate the bioenergetic profile of cells treated with PI5P. As shown in Figure 1D, mitochondrial respiration was dramatically reduced in cells treated with cell-permeant PI5P, with a significant reduction in basal and maximal respiration, as well as the spare respiratory capacity (Figure 1E). Importantly, PI5P-treated cells had a lower oxidative bioenergetic profile both in basal and stressed conditions, as indicated by the energy map (Figure 1F). We next analyzed the expression profile of key mitochondrial respiratory chain complexes (OXPHOS). While the total expression levels of NDUFB8 (representative of complex I) and SDHB (representative of complex II) were not altered in PI5P-treated cells, we found that the levels of UQCRC2, MTCO1 and APT5A (representatives of complexes III, IV and V) were upregulated in treated cells (Figure 1G). Such dysregulation of the OXPHOS stoichiometry, particularly between the electron transport chain (complexes I to IV) and ATP synthase (complex V) is indicative of a mitochondrial dysfunction. Taken together, these data indicate that PI5P treatment impairs mitochondrial function, preventing cardiomyocytes to cope with high energy demands.

**Figure 1.**
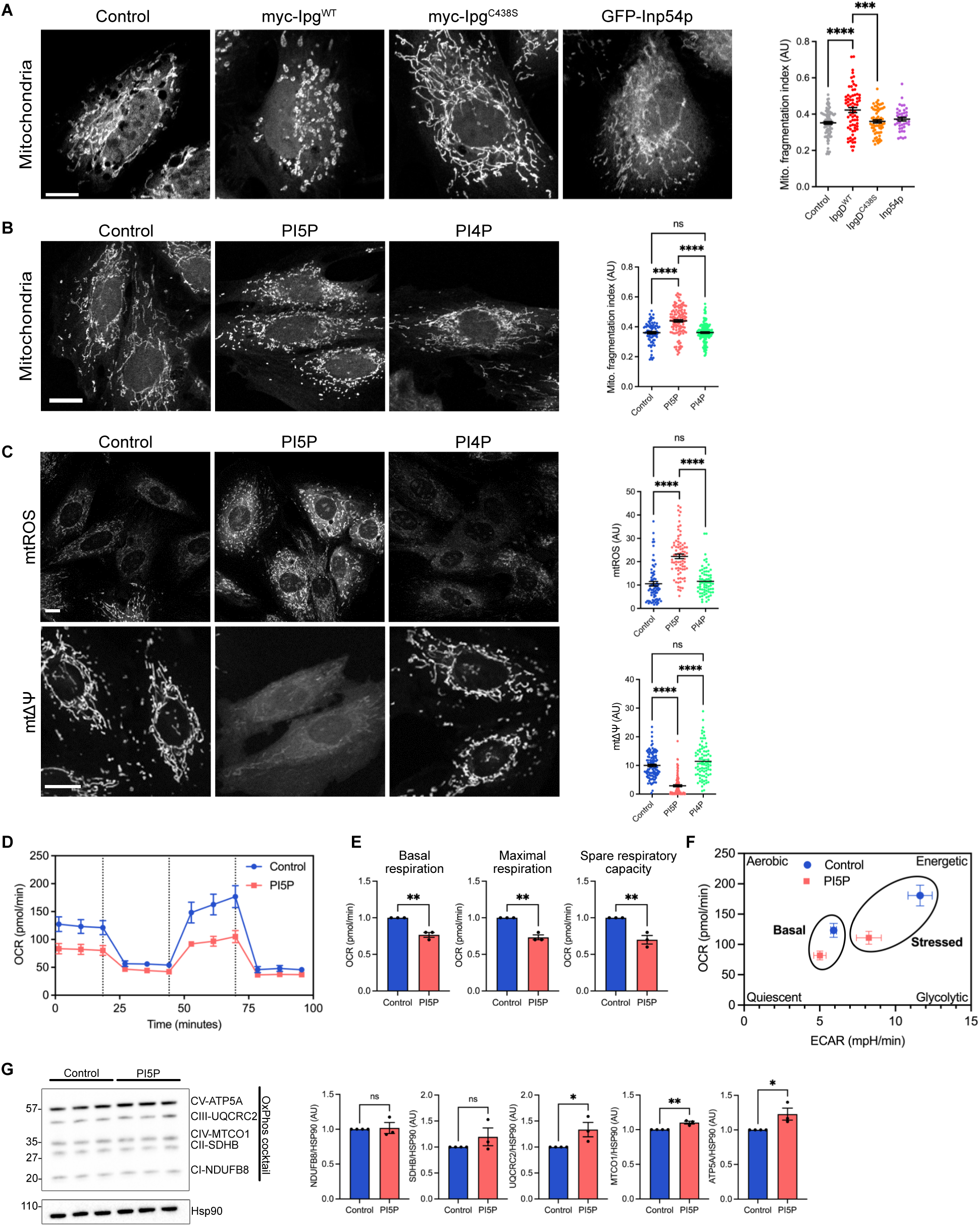
Exogenous PI5P triggers mitochondrial dysfunction. (A) Overexpression of IpgD induces mitochondrial fragmentation. Cells expressing myc-IpgD WT, C438S mutant or GFP-Inp54p were fixed and stained for mitochondria (left panel), and the mitochondrial fragmentation index was quantified across 3 independent experiments (right panel). Bar is 10µM in all figures unless otherwise stated. (B-C) Exogenous PI5P induces mitochondrial fragmentation and increases mitochondrial ROS (mtROS) production and decreased mitochondrial membrane potential (mtΔΨ). Cells were treated overnight with 15µM of PI5P or PI4P, fixed and stained for mitochondria (B) or incubated with MitoSOX (for mtROS production) or with TMRE (for mtΔΨ) and live-imaged (C). Quantifications from 3 independent experiments are shown on the right. (D) Bioenergetic profile of cells treated with PI5P. Treated cells were processed for a Mito Stress Test on a Seahorse flux analyser. A representative Seahorse trace including five technical replicates is shown. (E) Quantification of Basal respiration, Maximal respiration and Spare respiratory capacity from 3 independent experiments are shown. (F) Energy map from control or PI5P-treated cells showing a decrease in the oxidative capacity of the PI5P-treated cells, both in basal and stressed (FCCP treated) conditions. (G) Lysates from control or PI5P-treated cells were probed for the indicated antibodies (left panel). Quantifications are shown on the right. Statistical significance was determined by one-way ANOVA with Tukey’s post hoc analysis for multiple comparisons or unpaired t-test. *p<0.05, **p < 0.01, ***p < 0.001,****p < 0.0001, ns non-significant.

Next, we investigated whether the PI5P-induced mitochondrial dysfunction was impeding long-term cell viability. As shown in Figure S1A, PI5P-treatment did not alter cell viability as assessed by MTT staining, with similar viability in untreated or PI4P-treated cells even in long-term treatment or with high concentration. Moreover, no change in the expression levels of the pro-apoptotic markers Bax and p53 were detected (Figure S1B), neither in the expression level of the anti-apoptotic marker Bcl2 (Figure S1B).

### A significant pool of PI5P is localized at the mitochondria in cardiomyocytes

Given the effects of PI5P on mitochondria structure and function, we investigated in detail its subcellular localization using a previously described PI5P biosensor ^3,30,32^. As shown in Figure 2A and B, we observed two distinct localizations for endogenous PI5P: 1-on punctate vesicular structures, localized throughout the cell, with an accumulation in the juxtanuclear region, most likely representing an endosomal population as we previously described ^30,33^, and 2-a mitochondrial localization as highlighted by the colocalization with the mitochondrial marker Hsp60. The quantification of the signal intensity relative to the mitochondrial mask indicates that 24+/-1.5% of the endogenous PI5P is localized at the mitochondria (Figure 2C), revealing a mitochondrial pool of PI5P.

**Figure 2.**
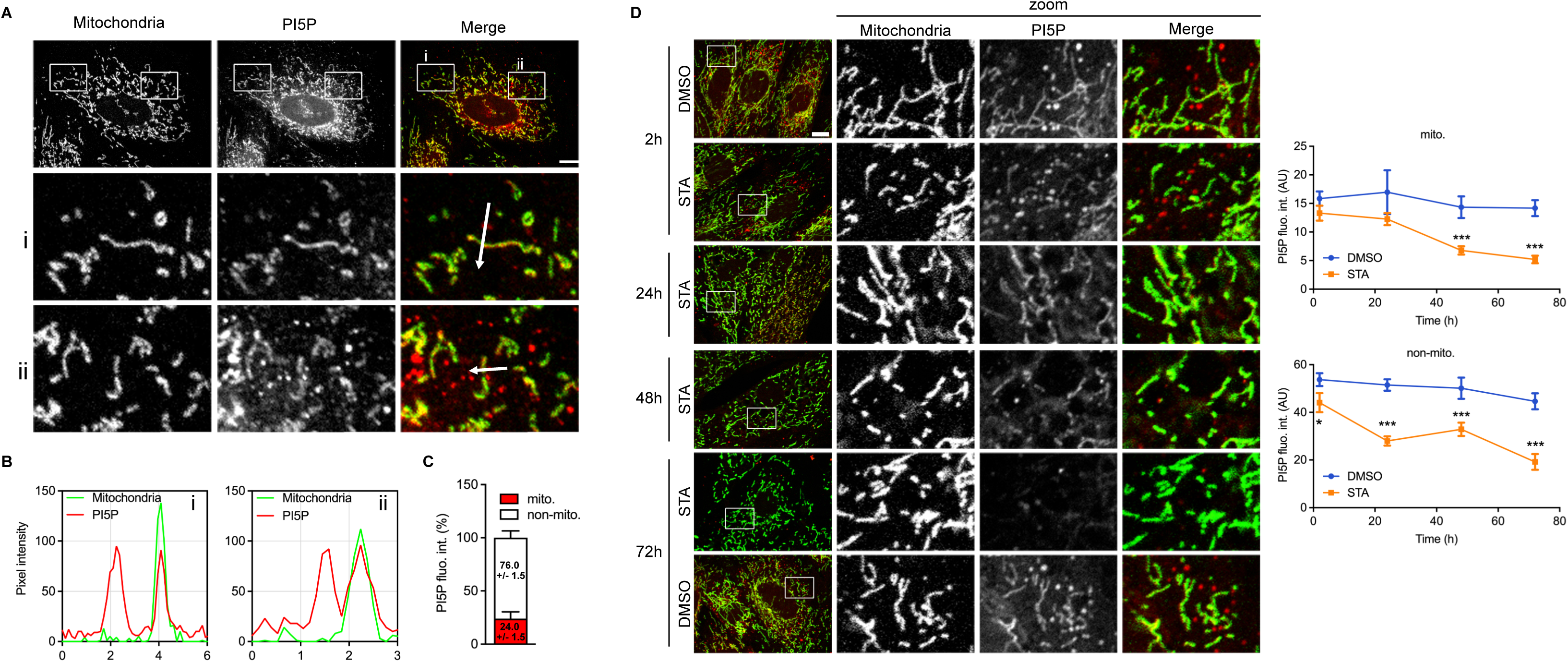
Subcellular localization of PI5P in cardiomyoblasts. (A) H9C2 cells were fixed and stained for mitochondria (in green) or for PI5P (in green). Zoomed region (i) and (ii) are shown below. (B) Line scan of fluorescence intensity along each arrow from inserts (i) and (ii) from (A). (C) Proportion of mitochondrial vs. non-mitochondrial PI5P. Results are from 19 cells across 3 independent experiments. (D) Cells were treated with STA (100nM for the indicated time) or with DMSO (vehicle only), fixed and stained for mitochondria (in green) or PI5P (in red). Right panel show the quantification of PI5P fluorescence intensity in the mitochondrial (mito.) or non-mitochondrial mask (non-mito.). Results are from 13-27 cells across 3 independent experiments. Statistical significance was determined by unpaired t-test. *p<0.05, ***p < 0.001.

### Both vesicular and mitochondrial PI5P pools are PIKfyve-dependent

To determine whether this mitochondrial PI5P pool was dependent on PIKfyve activity, we treated the cells with the selective PIKfyve inhibitor STA and quantified the different subcellular pools of PI5P in H9C2 cells. As shown in Figure 2D, in cells treated for 2h with STA, we observed a significant reduction of non-mitochondrial PI5P, as we described before in other cells types ^34^. Importantly, longer treatment further reduced the non-mitochondrial PI5P levels, down to 43+/-0.08% of the initial value after 72h of treatment. Interestingly, the mitochondrial pool of PI5P appeared also to be sensitive to PIKfyve inhibition, but with a different kinetics, being significantly reduced only after 48h of treatment. Longer treatment (ie 72h) failed to reduce further the levels of mitochondrial PI5P (down to 39+/-0.05% of the initial value). Altogether, these data indicate that both the non-mitochondrial and the mitochondrial pools of PI5P are dependent on PIKfyve kinase activity.

### Metabolic stress is associated with increased mitochondrial PI5P, possibly involving PIKfyve

Our previous work demonstrated that global cellular PI5P levels increased in a PIKfyve-dependent manner in cardiomyocyte, challenged by either hypoxic or metabolic stress ^27^. In order to investigate whether the mitochondrial pool of PI5P was also regulated during metabolic stress, we treated the cells with the glycolysis inhibitor 2-deoxy-D-glucose (2DG). 2DG-treatment induced a ∼2-fold increase in total PI5P levels (Figure 3A), consistent with what we previously observed using a mass-assay ^27^. Interestingly, 2DG-treatment increased both mitochondrial and non-mitochondrial PI5P levels (Figure 3A). To test if this stress-induced PI5P increased was also dependent on PIKfyve, we concomitantly treated the cells with STA. As shown in Figure 3B, and consistently with results obtained in Figure 2D, we found that STA-treatment prevented 2DG-induced PI5P increase for the non-mitochondrial pool. Regarding the PI5P mitochondrial pool, we found that PIKfyve inhibition by STA tended to decrease mitochondrial PI5P levels induced by 2DG after 4h of treatment. This is consistent with results obtained in basal conditions (Figure 2D), where we found that long-term STA-treatment are needed to decrease significantly mitochondrial PI5P. However, it has to be noted that long-term 2DG treatment causes apoptotic cell death of cardiomyoblasts ^27^, precluding us to investigate this further.

**Figure 3.**
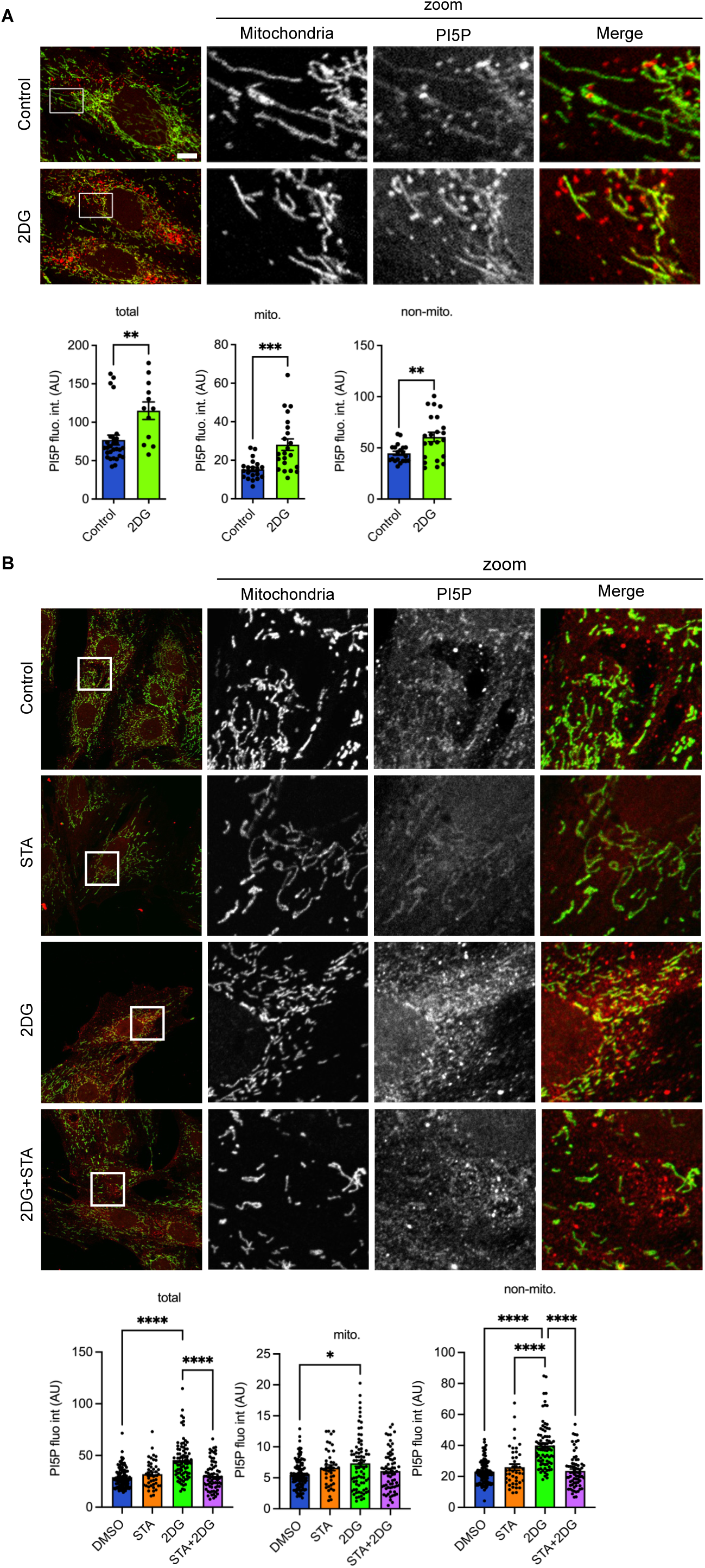
Metabolic stress increases mitochondrial and non-mitochondrial PI5P in a PIKfyve-dependent manner. (A) Cells were treated with 2DG (50mM) in complete medium for 4h, fixed and stained for mitochondria (in green) or PI5P (in red). Quantifications of total PI5P fluorescence, PI5P fluorescence intensity in the mitochondrial (mito.) or non-mitochondrial mask (non-mito.) from 3 independent experiments are shown on the bottom panel. (B) Cells were treated as in (A) except for a pre-treatment with STA (30min, 100nM) as indicated. Results are from 3 independent experiments. Statistical significance was determined by one-way ANOVA with Tukey’s post hoc analysis for multiple comparisons or unpaired t-test. *p<0.05, **p < 0.01, ***p < 0.001,****p < 0.0001.

### PIKfyve-endolysosomes are localized in close proximity to mitochondria

PIKfyve is known to be localized to the endolysosomal compartment, therefore the regulation by PIKfyve of the mitochondrial pool of PI5P raised questions. Therefore, we investigated the specific localization of endogenous PIKfyve in cardiomyocytes. PIKfyve presented a vesicular localization in cells, consistent with an endolysosomal localization, similarly to EEA1-staining. However, we found that several PIKfyve- or EEA1-positive vesicles showed a close proximity to the mitochondrial network (Figure 4A). To quantify the number of vesicles in immediate proximity with the mitochondria, we used a home-made analysis plugin for Image J as described in the methods section. As shown in Figure 4A, while we found that 20+/-1.6% of EEA1-positive vesicles were in close proximity with mitochondria, this number was significantly higher for PIKfyve-positive vesicles, up to 34+/-1.9%. Such a proximity prompted us to investigate the possible interaction of PIKfyve with cardiolipin, a major and specific mitochondrial lipid that accounts for 10–15% of the total mitochondrial phospholipids ^35^. PIKfyve immunoprecipitated from mouse heart (Figure S2A) was able to bind to PI3P, as previously described ^13^, but failed to interact with cardiolipin (Figure S2B), suggesting that the proximity of PIKfyve-endolysosomes with mitochondria is not mediated by an interaction with cardiolipin.

**Figure 4.**
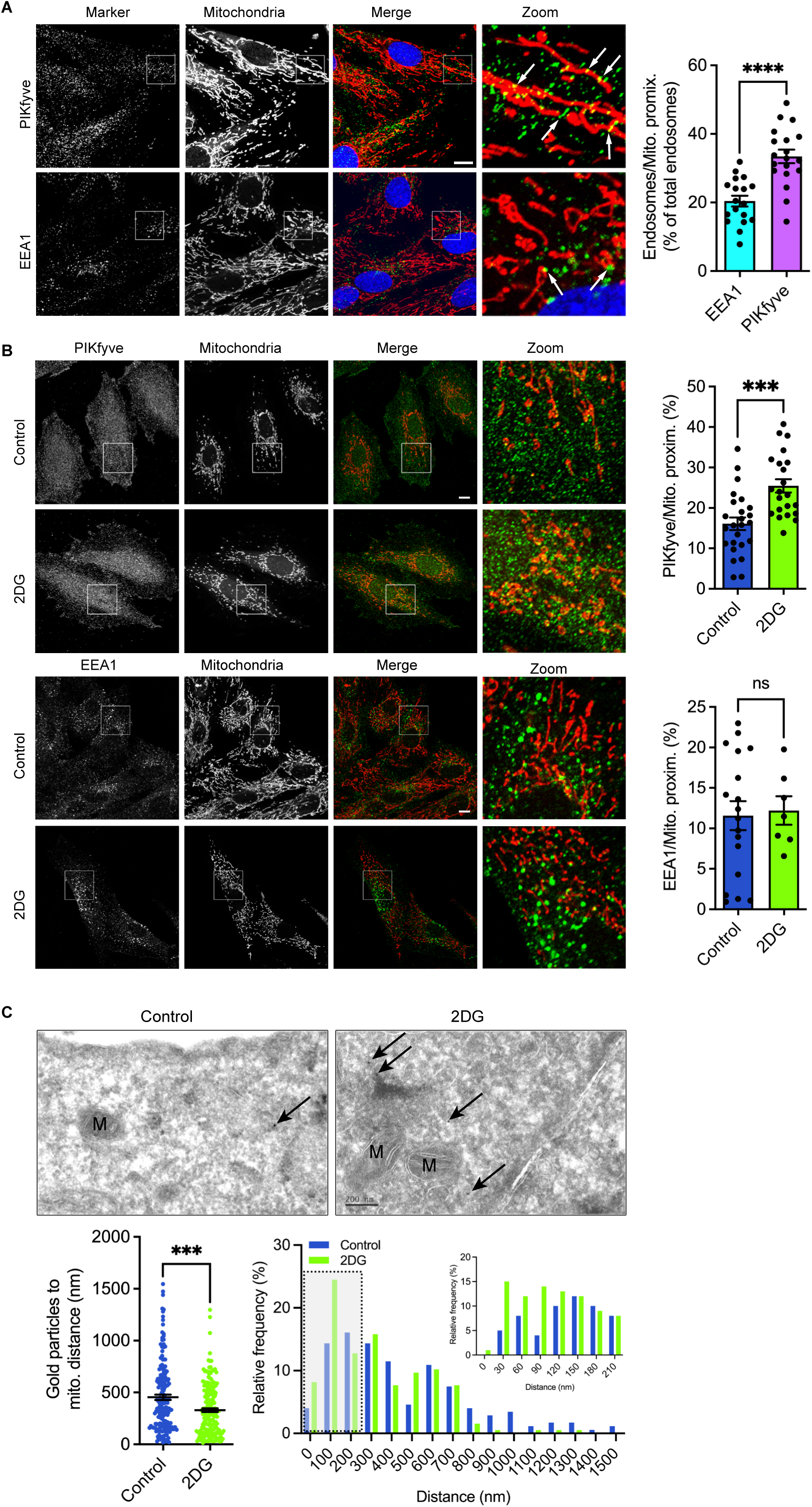
Metabolic stress increased PIKfyve-positive endolysosomes proximity with mitochondria. (A) Cells were fixed and stained for endogenous PIKfyve (upper panel, in green) or EEA1 (lower panel, in green) and with mitochondria (in red). Nuclei were stained with DAPI (in blue). Endosome to mitochondria proximity was assessed as described in the Methods section and is shown on the left. Results are from 3 independent experiments. (B) 2DG increases PIKfyve-but not EEA1-endosomes proximity with mitochondria. Results are from 3-4 independent experiments. (C) Immunogold labelling for endogenous PIKfyve in cells treated or not with 2DG. Left panel shows the quantification of the distance of gold particles to the closest mitochondria. Right panel show the relative frequency distribution of the measured distances. Inset depicts frequency distribution in the short-range distance (less than 210nm). Statistical significance was determined by unpaired t-test. ***p < 0.001,****p < 0.0001, ns non-significant.

### Metabolic stress increases the association of PIKfyve-endolysosomes with mitochondria

Next, we sought to investigate whether a metabolic stress was modifying this proximity. As shown in Figure 4B, 2DG-treatment increased significantly the PIKfyve-mitochondria proximity, while having no effect on EEA1-vesicles. This was further confirmed by immunogold labelling of cells, where we found a significant decrease in the measured distance between gold particles (ie endogenous PIKfyve) and mitochondria after 2DG-treatment (Figure 4C) and an increase in the frequency of particles in the short-range distance (<200nm). In order to confirm biochemically this proximity, we resorted to a subcellular fractionation protocol. Namely, a post-nuclear supernatant (PNS) obtained by controlled mechanical disruption of the cells was further centrifugated to generate a supernatant fraction (SPNT) and a pellet fraction (Figure 5A). Characterization of the obtained fractions is depicted in Figure 5B. We found that mitochondria were enriched in the pellet fraction (indicated by ATP5A), while the endoplasmic reticulum (ER, indicated by Calreticulin) and the early endosomes (EEA1) were equally distributed between the SPNT and the pellet. On the contrary, the cytosolic marker RhoGDI was excluded from the pellet fraction, similarly to the Golgi apparatus (Golgin-97). Interestingly, the late endosomal compartment, represented by Rab7, was marginally present in the pellet fraction, similarly to PIKfyve. Metabolic stress induced by 2DG did not alter the total expression of PIKfyve, as assessed either by qPCR (Figure S3A) or by western-blot in the PNS fraction (Figure S3B). However, we found that the amount of PIKfyve associated with the pellet fraction was significantly increased in 2DG-treated cells, together with Calreticulin (Figure 5B). This suggests an increased association between the mitochondria, the PIKfyve endolysosomes and the ER. Notably, no enrichment was found for either EEA1 nor Rab7, indicating that only the association of PIKfyve endolysosomes with mitochondria is regulated by metabolic stress.

**Figure 5.**
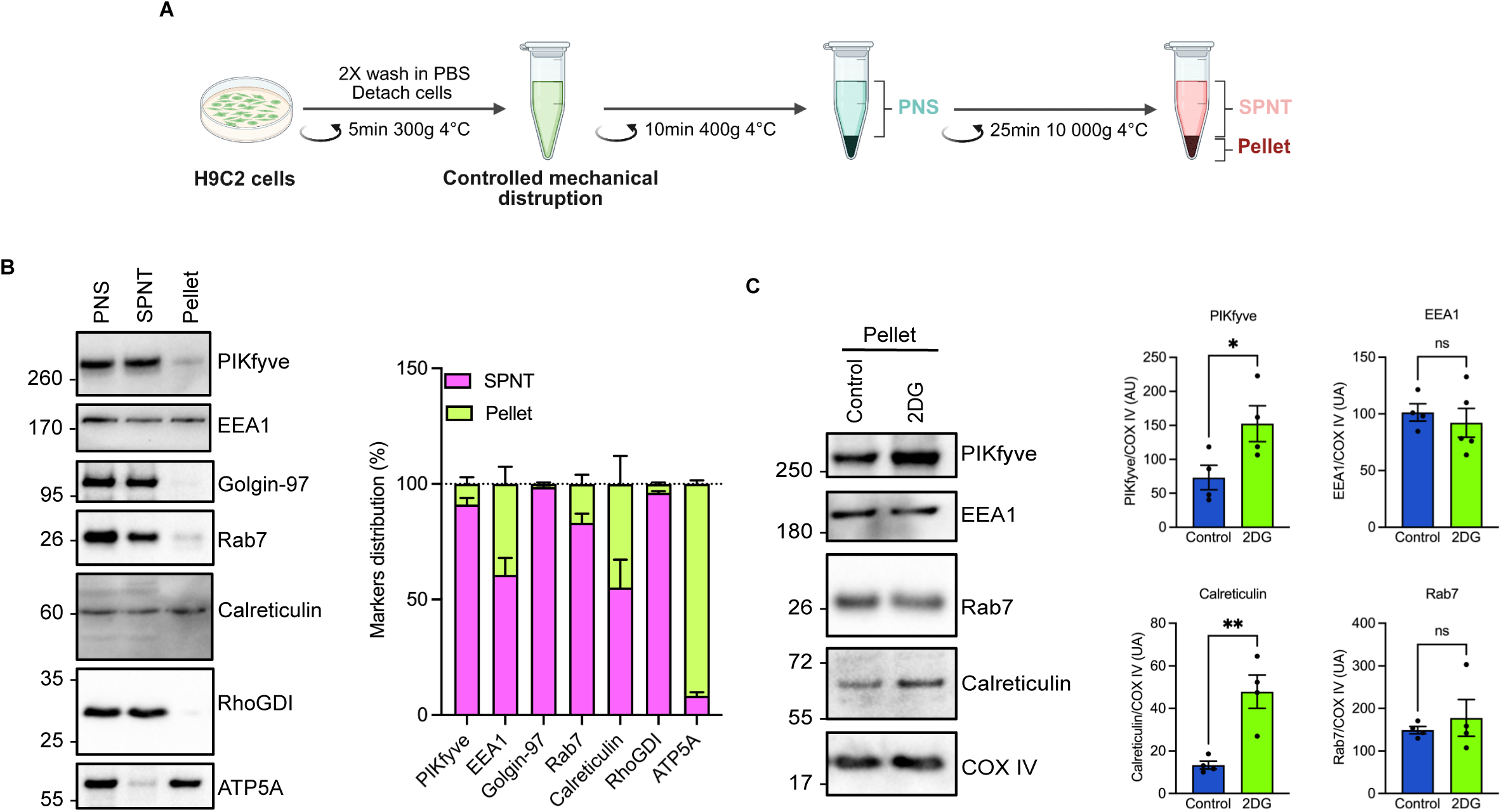
Biochemical association of PIKfyve with mitochondria is augmented by metabolic stress. (A) Schematic depicting the subcellular fractionation performed on H9C2 cells to obtained a post-nuclear supernatant (PNS), supernatant (SPNT) and pellet fractions. (B) The PNS, SPNT and pellet fractions were immunoblotted with the indicated antibodies. Quantification was performed from 4-8 independent experiments. (C) The pellet fractions obtained from control or 2DG-treated cells were immunoblotted with the indicated antibodies. Quantifications are shown on the right. Statistical significance was determined by unpaired t-test. *p<0.05, **p < 0.01, ns non-significant.

### PIKfyve-endolysosomes association with mitochondria is dependent on actin cytoskeleton

To better characterize the dynamics of PIKfyve on the endolysosomal membrane, we resorted to fluorescence recovery after photobleaching (FRAP), in which individual PIKfyve-positive vesicles were photobleached and the fluorescence recovery was monitored. As shown in Figure 6A, the mobile fraction of GFP-PIKfyve on endolysosomal membrane was estimated at 0.592+/-0.009 and the half-life time at 4.7s in control cells. In contrast, treated cells exhibited a lower mobile fraction (0.432+/- 0,008) with a higher half-life time (5.6s). Therefore, we concluded that GFP-PIKfyve exhibits long-term interaction with the endolysosomal membrane in 2DG-treated cells, as evidenced by the slower fluorescence recovery and the higher immobile fraction.

**Figure 6.**
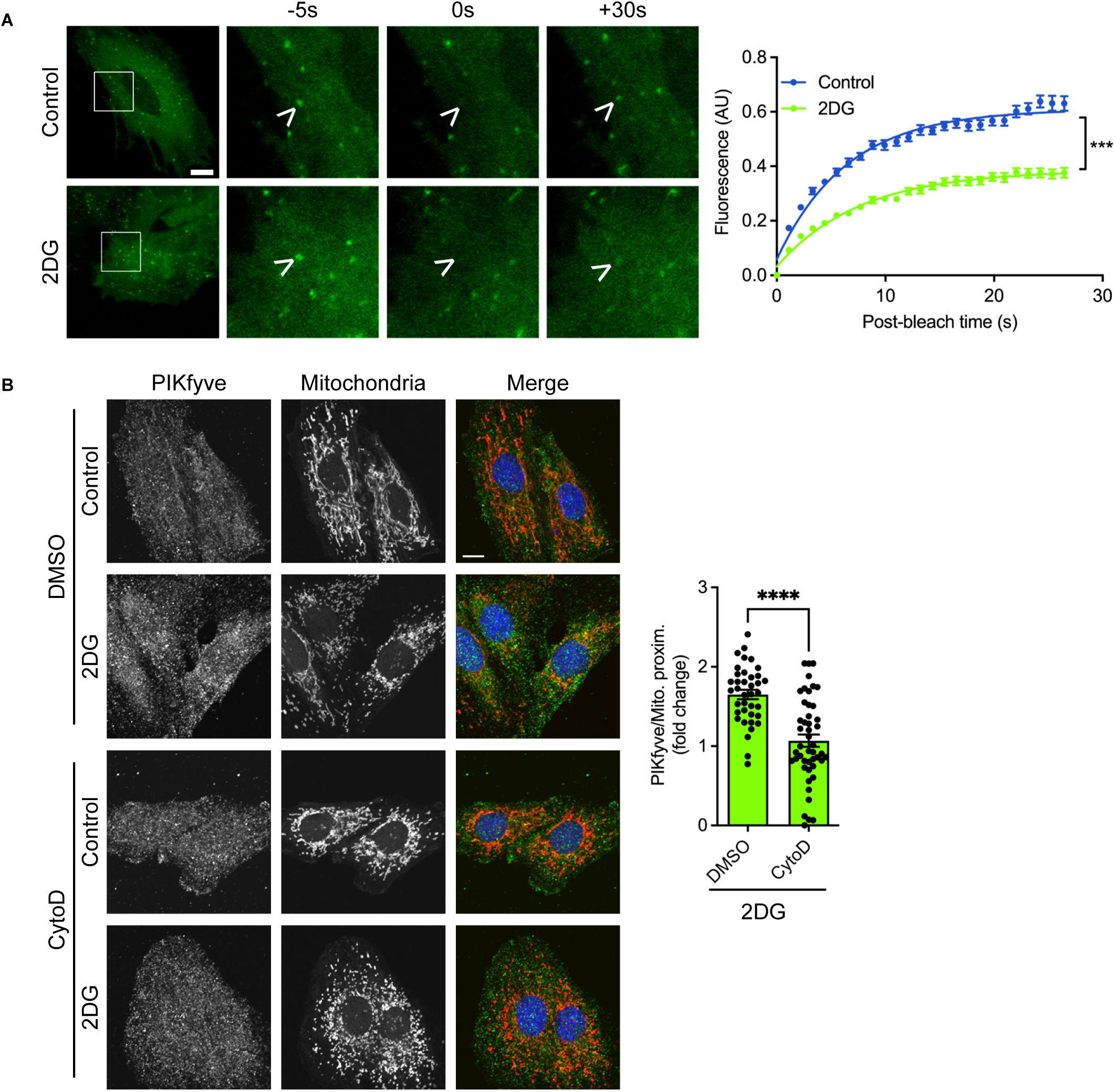
PIKfyve dynamics is impeded in stressed conditions dependently of actin mesh. (A) FRAP experiments was performed on GFP-PIKfyve overexpressing cells, in control or 2DG stimulated conditions. The fluorescence recovery after bleaching is shown on the right. Results are from 15-21 cells and 22 to 32 individual regions of interest across 4 independent experiments. (B) Cells were pre-treated for 30min with cytochalasin D (CytoD, 2µM) before being submitted to metabolic stress using 2DG (50mM, 4h), fixed and stained for PIKfyve (shown in green) and mitochondria (in red). Nuclei were stained with DAPI (in blue). PIKfyve to mitochondria proximity was quantified and is presented as fold change compared to control cells, from 3 independent experiments. Statistical significance was determined by unpaired t-test. ***p < 0.001,****p < 0.0001.

The actin cytoskeleton is known to be a key regulator of mitochondrial structure and function ^36^, and regulates several organelles interactions ^37^. Therefore, we next investigated whether actin cytoskeleton was involved in the PIKfyve-positive endolysosomes to mitochondria proximity. We found that Cytochalasin D, a potent inhibitor of actin polymerisation (as shown in Figure S4), prevented PIKfyve-endolysosomes to mitochondria proximity induced by 2DG (Figure 6B). Taken together, this suggests that actin filaments are needed to allow PIKfyve-endolysosomes and mitochondria proximity, presumably favouring PI5P transfer from the endolysosome membrane to the mitochondrial membrane.

## Discussion

In this study, we characterize a specific mitochondrial pool of PI5P in cardiomyocytes that contribute to mitochondrial dysfunction during a metabolic stress. We found that this PI5P pool relies on PIKfyve, the sole lipid kinase involved in PI5P synthesis in cells ^9,10^. Indeed, when PIKfyve is inhibited, both the vesicular and mitochondrial pools of PI5P are reduced, albeit with different kinetics: the vesicular pool is depleted first, followed by the mitochondrial pool. This differential turnover suggests that the primary site of PI5P synthesis is on the vesicles, most likely on endolysosomes as previously described ^30,34^, with some subsequent lipid transfer occurring to the mitochondrial membrane. Transfer of lipids between organelles, particularly phosphoinositides, has been extensively studied in the recent years (see ^38^ for review), specifically as it coordinates communication routes between organelles, and regulates their functions, and there is increasing evidence that membrane contact sites and lipid transfer between mitochondria and other organelles play a role in regulating mitochondrial function ^39,40^. For instance, it has been found that the Oxysterol binding protein related protein 1L (ORP1L) is involved in the transfer of PI4P from the late endosomal membrane to the mitochondria, a process implicating a three-way contact between the ER, the endolysosomes and the mitochondria and regulating mitochondrial fission ^41^. Moreover, PI3P has been proposed to regulate communication between the ER, early endosomes and mitochondria, allowing the regulation of ER structure, mitochondrial fission and ultimately controlling the energetic metabolism of the cell ^42^. The molecular machinery regulating these contact sites has been investigated. Wong and coworkers have found that the small GTPase Rab7, together with its Rab-GAP TBC1D15, regulate the positioning of the lysosomes close to the mitochondria, further regulating mitochondrial fission ^43^. Interestingly, this association is relevant in several pathological contexts, such as Charcot-Marie-Tooth type 2 disease ^44^, or acute myocardial infarction ^45^. In our case, we found that while PIKfyve-endolysosomes association with mitochondria was increased in metabolically-stressed cardiomyoblasts, Rab7 was not. This suggests that Rab7, and by extension TBC1D15 might not be involved in the establishment of PIKfyve-positive endolysosomes to mitochondria contacts, and hypothetically PI5P transfer to the mitochondrial membrane. More work is needed to characterize the molecular complex involved in the transfer of PI5P from the endolysosomes to the mitochondria.

Our work demonstrates that increased levels of PI5P induce a dysregulation of mitochondrial function, with the establishment of an unbalance between ETC and ATP synthase, resulting in an increased ROS production, decreased mitochondrial membrane potential and ultimately leading to energetic failure of the cell. It has been shown that pharmacological inhibition of PI5P4Kα/β, a 4-kinase responsible for the conversion of PI5P to PI(4,5)P_2_, induces a decrease in mitochondrial function, with impaired respiratory capacity ^46^. Similarly, the PI5P4Kα/β double knockout induces a reduced mitochondrial membrane potential and impaired mitochondrial respiration ^47^. Conversely, the depletion of PTPMT1, a PTEN-like 5-phosphatase with a strong preference for PI5P ^48^, induces an alteration of the mitochondrial aerobic metabolism ^49^. Interestingly, alterations of both PI5P4Kα/β and PTPMT1 are expected to induce an increase in PI5P levels, recapitulating the results we obtained in our study using cell-permeant lipid. Altogether, these data clearly suggest that high levels of PI5P are able to alter the cellular metabolism, inducing a mitochondrial dysfunction.

To date, the molecular effectors activated by PI5P in the mitochondria are still to be characterized. Yu and colleagues ^49^ have shown in vitro that the favored substrates of PTMT1 (namely PI(3,5)P_2_, PI(3,4)P_2_ and PI5P) are able to activate UCP2, therefore uncoupling the mitochondria and possibly explaining the energy deficiency observed in treated cells ^49^. Moreover, it has been shown that PTPMT1 regulates the phosphorylation of succinate dehydrogenase, therefore regulating the energetic status of the cell ^50^. However, the specific implication of the phosphoinositides regulated by PTPMT1 were not characterized in this context. Further work is needed to identify the molecular effectors of the mitochondrial PI5P.

In conclusion, our work has revealed an unexpected role for PIKfyve in regulating the dialogue between the endolysosomal and mitochondrial systems, and has identified a PIKfyve-dependent mitochondrial pool of PI5P that contributes to mitochondrial dysfunction and metabolic abnormalities.

## Methods

### Reagents and antibodies

Antibodies used in this study are as follows: anti-myc (9E10, sc-40), anti-Hsp60 (H1, sc-13115), anti-PIKfyve (64-Q6, sc-100408 for immunofluorescence) and anti-RhoGDI (sc-373723) from Santa Cruz Biotechnology, anti-PIKfyve (A6689, Abclonal for western blot), anti-EEA1 (610457 clone 14, BD Biosciences), anti-Golgin-97 (CDF4, eBioscience), anti-Rab7 (D95F2 XP, Cell Signaling Technology), anti-Calreticulin (PA3-900, ThermoFisher Scientific), anti-ATP5A (ab110413, Abcam), anti-COX IV (ARG66326, from Arigo Biolaboratories). Fluorescent Alexa-coupled secondary antibodies and DAPI were from Life Technologies and HRP-coupled secondary antibodies from Cell Signaling Technology. STA was purchased from Axon Medchem. Plasmids used in this study were described before: pcDNA3-GFP-PIKfyve ^7^, pRK5-myc-IpgD, prK5-myc-IpgD^C438S^ and peGFP-Inp54p ^30^. All other chemicals were from Sigma unless otherwise stated.

### Cell culture, transfection and treatments

The rat embryonic cardiomyoblastic cell line H9C2 (2-1, Merck) was cultured in DMEM medium (31966, ThermoFisher Scientific) supplemented with 10% FBS and 1% penicillin– streptomycin in a 37°C, 5% CO2 incubator. Cells were transfected either with TransIT®2020, TransIT-X2 or X-tremeGENE9 (Merck) according to manufacturer’s instructions. To induce metabolic stress, the cells were treated with 2-deoxy-D-glucose (2DG, 50mM) in complete medium for 4h, as described before ^27^. Cells were treated with STA (100nM), DSMO (vehicle only) or cell permeant di-C4 PI5P (P-5004, Echelon Biosciences) or PI4P (P-4004, Echelon Biosciences) as indicated. Cell viability was assessed with MTT colorimetric assay.

### Immunofluorescence and live-cell imaging

For immunofluorescence, cells were grown on glass coverslips, fixed with PFA, processed as described ^51^ and imaged on a Zeiss LSM-780 or LSM-900 confocal microscope. The quantification of endosomes-mitochondria proximity was performed in ImageJ. Briefly, the images were thresholded, and the Boolean operator AND was used to define a proximity mask between the two channels. Using the ROI manager, the number of detected particles in the proximity mask was normalized against the total number of particles detected per cell and expressed as percentage. For PI5P detection, we used the previously described biotinylated PHD probe in combination with a streptavidin-Alexa564 ^34^. FRAP experiments were performed as before ^52^. Essentially, cells were seeded on glass-bottomed dishes and were treated 24h post-transfection in complete medium with 2DG (50mM, 4h). The culture medium was then changed to imaging medium (HBSS+20mM HEPES) and live-imaged in a 5% CO_2_ 37°C heated chamber on a LSM780 microscope piloted with Zen Software.

### Immunoelectron Microscopy

For immunoelectron microscopy, samples were fixed with a mixture of 2% PFA and 0.2% glutaraldehyde in 0.1 M phosphate buffer (pH 7.4). Cells were scraped, pelleted, and embedded in 10% bovine skin gelatin in 0.1 M phosphate buffer. Fragments of the pellet were infiltrated overnight with 2.3 M sucrose in 0.1 M phosphate buffer at 4°C, mounted on aluminum studs, and frozen in liquid nitrogen. Sectioning was done at −120°C in a cryo-ultramicrotome (UC7/FC7 Leica Microsystems). The 80-nm ultrathin sections were collected in 1:1 mixture of 2.3 M sucrose and 2% methyl cellulose and transferred onto Formvar-carbon-coated nickel grids for immunogold localization. Cells were immunolabeled with an anti-PIKfyve antibody (20µg/mL, sc-100408, Santa Cruz) followed by a bridging antibody (10µg/mL, 610-4120, Rockland) and finally immunogold-labeled using 10 nm protein A-gold particles. Grids were then stained with a mixture of 2% methylcellulose and 0.4% uranyl acetate. Samples were analyzed with a TEM (Jeol JEM-1400, JEOL) at 80 kV. Images were acquired using a digital camera (Gatan Orius, Gatan) at 10,000× magnification.

### Subcellular fractionation, protein extraction and western blotting

Subcellular fractionation was performed as described ^27^. Briefly, cells were washed twice in PBS, detach using Trypsin-EDTA, centrifugated 5min at 300g 4°C. The cell pellet was resuspended in HB (20mM HEPES pH 7.4, 270mM de sucrose, proteases inhibitors (P8340, Sigma) and phosphatases inhibitors (78420, ThermoFisher Scientific) and disrupted by 10 passages through a 27G needle. The homogenate was centrifugated (10min 400g 4°C) and the obtained post-nuclear supernatant (PNS) was further centrifugated 25min 10000g 4°C to generate a supernatant fraction and a pellet fraction. The pellet was gently rinsed in HB and solubilized in complete RIPA buffer. Proteins were quantified using the BCA Protein Assay (B9643 and C2284, Sigma-Aldrich). Proteins were loaded in Laemmli sample buffer, denaturated at 70°C for 15min or at 37°C for 30min (for ATP5A antibody), and resolved by SDS–PAGE and Western blotting. Immunoreactive bands were detected by chemiluminescence with the Clarity Western ECL Substrate (Bio-Rad) on a ChemiDoc MP Acquisition system (Bio-Rad). Quantification of immunoreactive bands was performed using ImageLab v6.1.0 (Bio-Rad) on non-saturated acquisitions.

### Immunoprecipitation and lipid-protein overlay assay

PIKfyve was immunoprecipitated from mouse heart as follows: Twenty mg of mouse heart were lysed in complete RIPA buffer in a FastsPrep-24 bead grinder, using Lysing matrix S. Lysates were clarified by centrifugation 20min 10000g 4°C, proteins were quantified using BCA and 2.5mg total proteins were further processed for immunoprecipitation using 20µl of anti-PIKfyve antibody (sc-100408), 150µl of a 50% slurry Protein G-Sepharose (Merck) in a final volume of 500µl. After o/n incubation at 4°C, the beads were extensively washed and further equilibrated in 10mM Tris-HCl pH7.5. Proteins were eluted by adding 100µl Glycine 0.1M pH2.5 for 10min, neutralized by adding 50µl 10mM Tris-HCl pH7.5. Elution was repeated and both elutions were pooled. The amount of eluted PIKfyve was estimated by western-blot against a range of total lysate, and further used for lipid-protein overlay assay. Lipids were as follows: PI3P diC16 (P-3016, Echelon Biosciences), 1-palmitoyl-2-oleoyl-glycero-3-phosphocholine (PC, 850457) and Cardiolipid (840012) from Avanti Polar. Lipids were diluted in a mixture of chloroform:methanol (1:1) and spotted on a Hybond C-extra nitrocellulose membrane (Amersham). The membrane was hydrated in TBS containing 0.1% Tween 20 (TBST), and the lipid-protein overlay was further performed as described ^4^, using approximately 150µg/ml immunoprecipitated PIKfyve according total lysates. Bound PIKfyve was detected using an anti-PIKfyve antibody (sc-100408) with an anti-mouse HRP-coupled.

### RNA Extraction and Quantitative Polymerase Chain Reaction (qRT-PCR)

RNA extraction and qRT-PCR were performed as before ^27,34^. Briefly, total RNAs were isolated from cells using the RNeasy mini kit (Qiagen), reverse transcribed using Superscript II reverse transcriptase (Invitrogen) in the presence of a random hexamers and processed for real-time quantitative PCR using MesaBlue qPCR MasterMix Plus (Eurogentec). The expression of target mRNA was normalized against house-keeping genes (GAPDH or RPLPO) mRNA expression. Primers used in this study are detailed in Table 1.

**Table 1.**
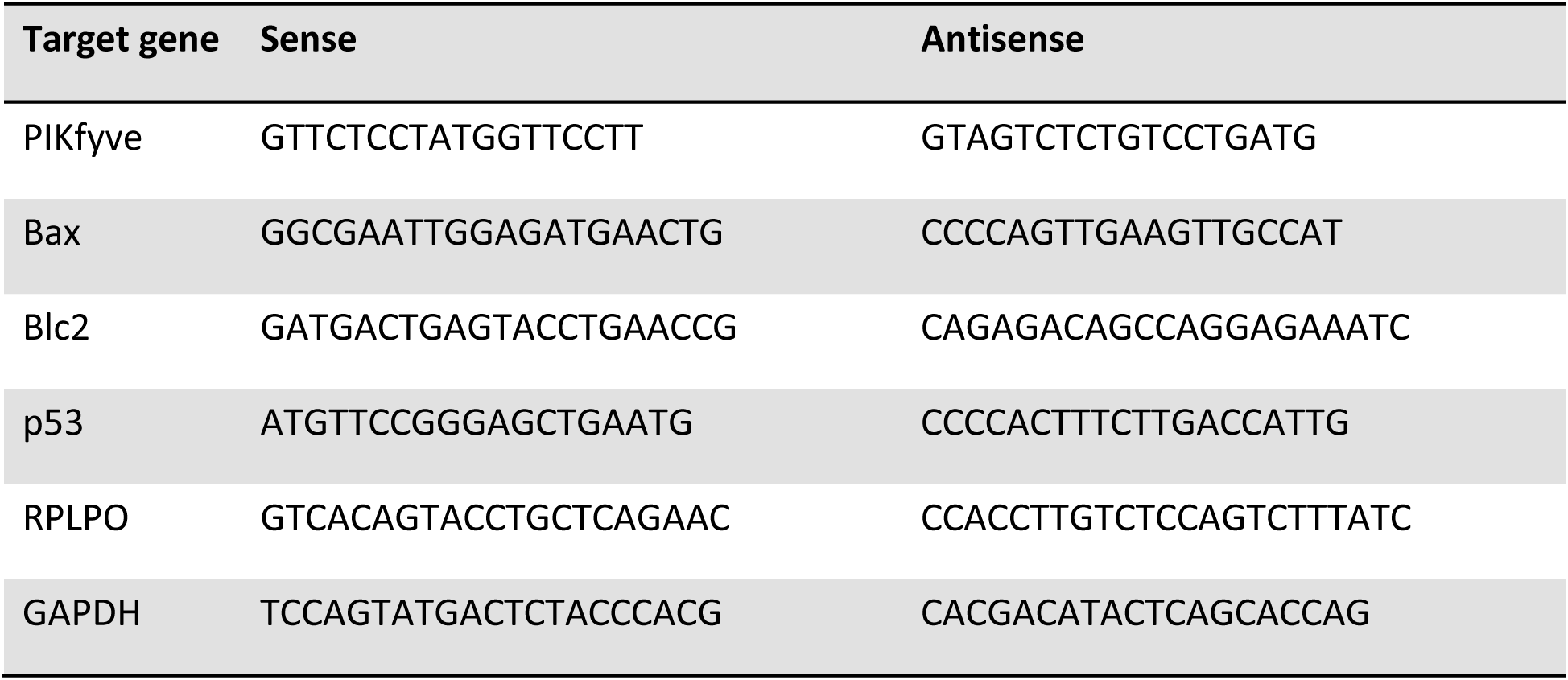
qRT-PCR primers used in the study.

### Metabolic profiling of cells

Bioenergetic profiling of cells was performed on a Seahorse flux analyzer XFe24. Briefly, cells were seeded on Seahorse XFe24 plates at the density of 75000 cells per well and after 4h attachment were treated in complete medium with cell-permeant PI5P overnight at 15µM. Cells were then washed twice in Seahorse XF DMEM supplemented with 10mM glucose, 4mM glutamine and 1mM sodium pyruvate, and equilibrated 1h in a CO2-free 37°C incubator. The Mito Stress test was performed using 1µM oligomycin, 4µM FCCP and antimycin A + rotenone (1µM each). Oxygen consumption rate (OCR) and extracellular acidification rate (ECAR) were monitored during sequential incubations and parameters were calculated according to Seahorse XF Cell Mito Stress Test Report Generator. The energy map was built according to the Seahorse XF Cell Energy Phenotype Test Report Generator under basal and FCCP-stressed conditions.

### Statistical analysis

Data are expressed as mean ± SEM. Comparison between two groups was performed by Student’s one-tailed t-test while comparison of multiple groups was performed by one-way ANOVA followed by a Tukey’s post hoc test using GraphPad Prism version 10.1.2 (GraphPad Software, LLC).

## Acknowledgments

We thank the Rangueil imaging platform (Genotoul-TRI) at the I2MC. We acknowledge the METi imaging facility (Genotoul-TRI), member of the national infrastructure France-BioImaging supported by the French National Research Agency (ANR-24-INBS-0005 FBI BIOGEN).

This work was funded by INSERM, Université de Toulouse, Fédération Française de Cardiologie (Dotation 2019), and Fondation de France (WB-2022-44948).

## Authors contributions

HT and FB designed and conceptualized the study. HT, SBH, MC, VA and FB performed the biochemical and cell biological experiments. SB and VS performed and analysed the electron microscopy experiments. HT and FB co-wrote the manuscript, with critical review and commentary from OK and FL.

