## Supplemental figures for "The phosphoinositide PI5P impairs mitochondrial function through endosome-mitochondria proximity"

**Figure S1. Long-term exposure to PI5P does not alter cell viability.**

(A) H9C2 cells were treated as indicated and cell viability was assessed by MTT staining. Results are shown as mean+/-SEM and are from 4 independent experiments performed in quadruplicate.

(B) qPCR analysis of indicated genes from cells treated or not with PI5P (15µM, 24h).

ns, non-significant from unpaired t test.

**Figure S2. PIKfyve does not interact with cardiolipin.**

(A) Endogenous PIKfyve was immunoprecipitated from two mouse hearts. Total heart lysate and elution fractions were immunoblotted for PIKfyve.

(B) Elution from Heart #1 in (A) was used on a lipid-protein overlay assay, showing the interaction of PIKfyve with PI3P but not with cardiolipin or phosphatidylcholine.

**Figure S3. Total expression levels of tested markers are unaffected by 2DG.**

(A) qPCR analysis of PIKfyve expression level in H9C2 cells treated or not with 2DG for 4h.

(B) Western-blot analysis of expression levels of indicated markers in control or 2DG-treated cells.

ns, non-significant from unpaired t test.

**Figure S4. Effect of cytochalasin D on actin cytoskeleton.**

H9C2 cells were treated or not with cytochalasin D, fixed and actin cytoskeleton was stained using fluorescently-labelled phalloidin (shown in green). Nuclei were stained with DAPI (in blue). Bar is 10µm.

**Figure S1**

A

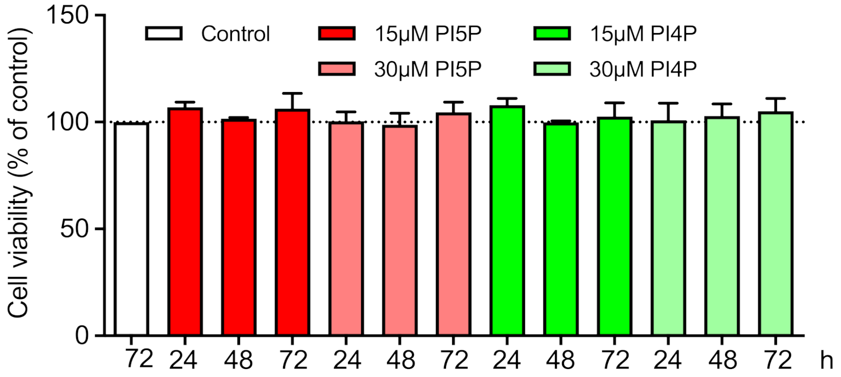
Cell viability (% of control)
150
100
50
0
Control
15μM PI5P
30μM PI5P
15μM PI4P
30μM PI4P
72 24 48 72 24 48 72 24 48 72 24 48 72 h

B

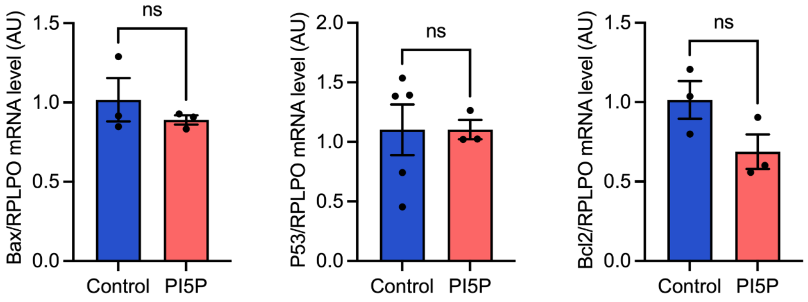
Bax/RPLPO mRNA level (AU)
1.5
1.0
0.5
0.0
ns
Control PI5P
P53/RPLPO mRNA level (AU)
2.0
1.5
1.0
0.5
0.0
ns
Control PI5P
Bcl2/RPLPO mRNA level (AU)
1.5
1.0
0.5
0.0
ns
Control PI5P

**Figure S2**

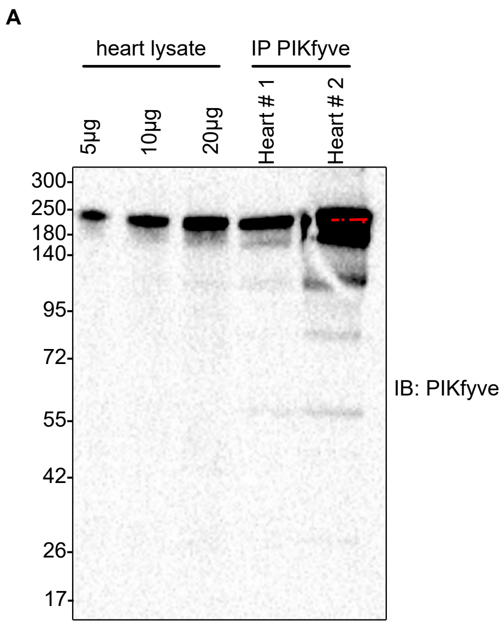
A
heart lysate
IP PIKfyve
5µg
10µg
20µg
Heart # 1
Heart # 2
300
250
180
140
95
72
55
42
26
17
IB: PIKfyve

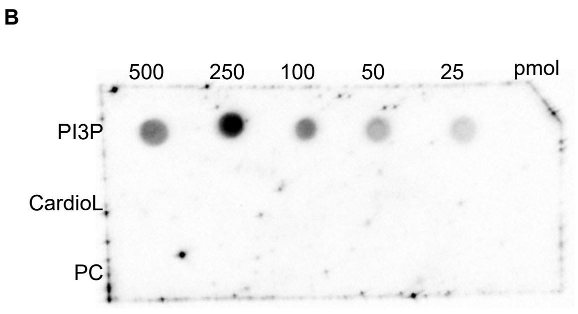
B
500
250
100
50
25
pmol
PI3P
CardioL
PC

**Figure S3**

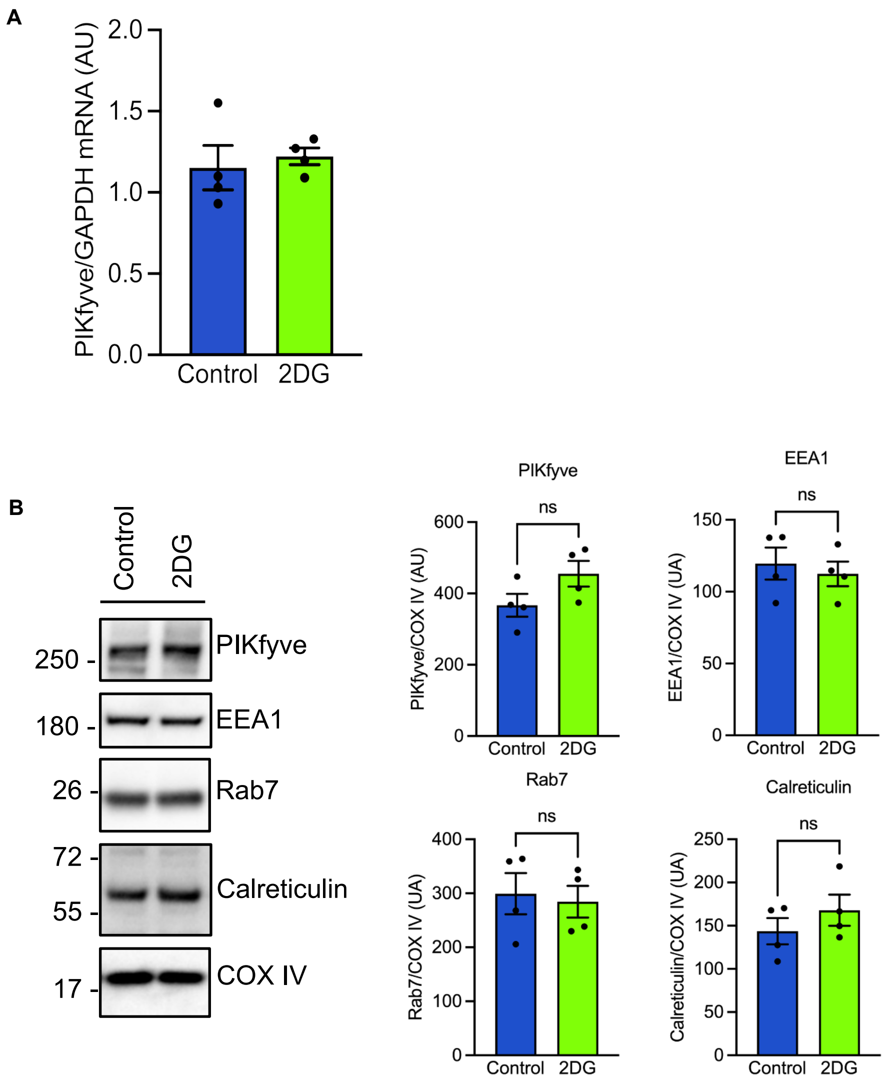

A
PIKfyve/GAPDH mRNA (AU)
2.0
1.5
1.0
0.5
0.0
Control
2DG
B
Control
2DG
250 -
PIKfyve
180 -
EEA1
26 -
Rab7
72 -
55 -
Calreticulin
17 -
COX IV
PIKfyve
ns
PIKfyve/COX IV (AU)
600
400
200
0
Control
2DG
EEA1
ns
EEA1/COX IV (UA)
150
100
50
0
Control
2DG
Rab7
ns
Rab7/COX IV (UA)
400
300
200
100
0
Control
2DG
Calreticulin
ns
Calreticulin/COX IV (UA)
250
200
150
100
50
0
Control
2DG

**Figure S4**

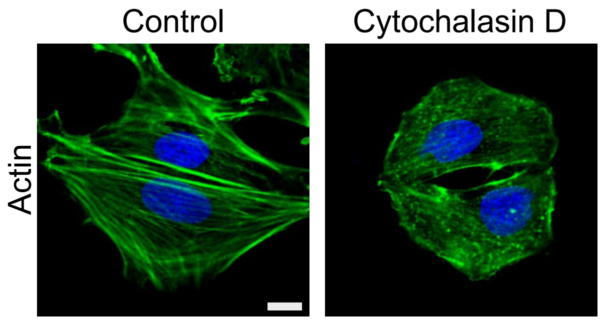
